# Distinct roles of *Bdnf I* and *Bdnf IV* transcript variant expression in hippocampal neurons

**DOI:** 10.1101/2023.04.05.535694

**Authors:** Svitlana V. Bach, Allison J. Bauman, Darya Hosein, Jennifer J. Tuscher, Lara Ianov, Kelsey M. Greathouse, Benjamin W. Henderson, Jeremy H. Herskowitz, Keri Martinowich, Jeremy J. Day

## Abstract

Brain-derived neurotrophic factor (*Bdnf*) plays a critical role in brain development, dendritic growth, synaptic plasticity, as well as learning and memory. The rodent *Bdnf* gene contains nine 5′ non-coding exons (*I-IXa*), which are spliced to a common 3′ coding exon (*IX*). Transcription of individual *Bdnf* variants, which all encode the same BDNF protein, is initiated at unique promoters upstream of each non-coding exon, enabling precise spatiotemporal and activity-dependent regulation of *Bdnf* expression. Although prior evidence suggests that *Bdnf* transcripts containing exon *I* (*Bdnf I*) or exon *IV* (*Bdnf IV*) are uniquely regulated by neuronal activity, the functional significance of different *Bdnf* transcript variants remains unclear. To investigate functional roles of activity-dependent *Bdnf I* and *IV* transcripts, we used a CRISPR activation (CRISPRa) system in which catalytically-dead Cas9 (dCas9) fused to a transcriptional activator (VPR) is targeted to individual *Bdnf* promoters with single guide RNAs (sgRNAs), resulting in transcript-specific *Bdnf* upregulation. *Bdnf I* upregulation is associated with gene expression changes linked to dendritic growth, while *Bdnf IV* upregulation is associated with genes that regulate protein catabolism. Upregulation of *Bdnf I*, but not *Bdnf IV*, increased mushroom spine density, volume, length, and head diameter, and also produced more complex dendritic arbors in cultured rat hippocampal neurons. In contrast, upregulation of *Bdnf IV*, but not *Bdnf I*, in the rat hippocampus attenuated contextual fear expression. Our data suggest that while *Bdnf I* and *IV* are both activity-dependent, BDNF produced from these promoters may serve unique cellular, synaptic, and behavioral functions.

## INTRODUCTION

Brain-derived neurotrophic factor (BDNF) is a major regulator of nervous system function. Signaling through the TrkB receptor, BDNF facilitates neuronal differentiation, proliferation, and survival, as well as axonal and dendritic growth, synapse formation, and maturation [1–3]. BDNF is critical for various forms of synaptic plasticity [4–7], and is implicated in many behavioral functions, including multiple types of learning and memory. A significant challenge in studying BDNF function is the complex regulation of the *Bdnf* gene. *Bdnf* consists of nine 5′ non-coding exons, designated as *I* – *IXa*, each containing a unique promoter from which transcription is initiated [8, 9]. These non-coding exons are spliced to a 3′ common-coding exon (*IX*) [10], such that all *Bdnf* transcript variants produce the same BDNF protein.

Transcriptional and epigenetic regulation at *Bdnf* promoters tightly controls temporal, spatial, and activity-dependent *Bdnf* expression. Exposure to a wide variety of stimuli, which engage different epigenetic elements and transcription factors, drives transcription of individual *Bdnf* variants from their upstream promoters [11, 12]. For example, *cis*-elements within *Bdnf* promoter *I* facilitate AP-1-dependent transcription [11] while dynamic changes in DNA methylation control transcription from *Bdnf* promoter *IV* following defeat stress and antidepressant treatment [12]. Transcript-specific regulation of *Bdnf* occurs across multiple rodent models and in response to seizures [13, 14], ischemia [15], and stress [16]. Moreover, production of multiple *Bdnf* transcripts contributes to BDNF regulation by controlling its activity-dependent and brain region-specific expression, subcellular localization patterns, and its local translation and transcript stability [17, 18].

Despite the known functional roles of BDNF protein, how individual *Bdnf* transcripts differentially contribute to BDNF-dependent cellular, circuit, and behavioral functions remains unclear. To address this question, we established a transcript-specific CRISPR activation (CRISPRa) system capable of selectively upregulating *Bdnf* variants *I* and *IV* from their endogenous promoters using a catalytically-dead Cas9 (dCas9) protein fused to a strong tripartite transcriptional activator, VPR, comprised of VP64-p65-Rta [19, 20]. While both *Bdnf I* and *IV* promoters are responsive to neuronal activity [8, 13], their activation kinetics differ, and they respond to different transcriptional and epigenetic signaling machinery. Here, we used our recently developed CRISPRa system to better understand the functional significance and individual contribution of *Bdnf* transcript *I* versus *IV* on BDNF-dependent functions. We show that *Bdnf I* upregulation in hippocampal neurons was associated with gene expression changes responsible for dendritic growth, while *Bdnf IV* upregulation was associated with protein catabolism. Upregulation of *Bdnf I*, but not *IV*, caused changes in dendritic spine dynamics in hippocampal neurons, while upregulation of *Bdnf IV* (but not *I*) in the rat hippocampus impacted contextual fear expression. These data indicate that although *Bdnf I* and *IV* transcripts encode the same protein, selective overexpression of these transcripts causes distinct downstream effects on gene expression, cellular morphology, and behavior.

## MATERIALS AND METHODS

### Animals and cell culture

All experiments were approved by the University of Alabama at Birmingham Institutional Animal Care and Use Committee (IACUC). Male Sprague Dawley rats (90-to 120-day old, 250-350 g) were co-housed in pairs on a 12/12 h light/dark cycle with *ad libitum* food and water. Timed pregnant dams were individually housed until embryonic day 18 for hippocampal culture harvest [20, 21]. Cells were maintained in complete Neurobasal media for 11 - 14 d *in vitro* (DIV) with half media changes at DIV1, 4–5, 8–9, and 12.

### CRISPR construct design and delivery

CRISRP/dCas9 and sgRNA construct design was previously described and all constructs are available on Addgene [20, 22]. Lentiviruses were produced in a sterile, BSL-2 environment [22]. Physical viral titer was determined using Lenti-X RT-qPCR Titration kit (Takara), and only viruses > 1 × 10^10^ GC/ml or > 1 × 10^12^ GC/ml were used for cell culture or *in vivo* experiments, respectively. Stereotaxic surgery was performed as previously described [20]. Bilateral infusions of 1.5 μl (0.5 μl sgRNA and 1 μl dCas9-VPR viruses in sterile 1X PBS, each hemisphere) were directed to dorsal CA1 (AP: –3.3 mm, ML: ±2.0 mm, DV -3.1 mm from bregma). Additional details can be found in supplemental methods.

### Spine morphology imaging and analysis

Automated image analysis was performed with Neurolucida 360 (2.70.1, MBF Biosciences, Williston, Vermont), as previously described [23]. After deconvolution, image stacks were imported into Neurolucida 360, the full dendrite length was traced with semi-automatic directional kernel algorithm. A blinded experimenter manually confirmed all assigned points and made any necessary adjustments. Each dendritic protrusion was automatically classified as a dendritic filopodium, thin spine, stubby spine, or mushroom spine [24]. Reconstructions of branched structure analysis were collected in Neurolucida Explorer (2.70.1, MBF Biosciences, Williston, Vermont). Spine density was calculated as the number of spines per 10 μm of dendrite length.

### Behavior

Contextual fear conditioning (CFC) was conducted as described [25]. Rats were placed into the training chamber with a metal floor grid (Med Associates) and allowed to explore for 7 min, during which three electric shocks (1 s, 0.5 mA each) were administered every 2 min. Memory was tested at 1 h, 24 h, and 7 days after training. An open field arena (43 × 43 cm; Med Associates) was used to assess locomotor and anxiety behavior, as previously described [25]. One week following CFC, rats were placed in the open field arena and allowed to explore for 30 min. Distance traveled (in cm) and time spent in the center (s) were tracked quantified using automated video tracking software (CinePlex Studio, Plexon Inc).

Additional details for all procedures can be found in supplemental methods.

## RESULTS

### Transcription of *Bdnf* variants I and IV is regulated by activity

*Bdnf* promoters respond differently to neuronal activity [10, 11, 13, 18]. Therefore, we first examined activity-dependent *Bdnf* transcript expression patterns in response to neuronal stimulation. DIV 11 rat hippocampal primary neuronal cultures were depolarized with 25 mM potassium chloride (KCl) for 1 - 4 hours and *Bdnf* transcripts were measured with RT-qPCR (**Fig. 1a-b**; primer sequences in **Table S1**). Expression of *Bdnf I, II, IV*, and *VI*, as well as total *Bdnf IX*, was altered by KCl depolarization, with peak changes occurring 3 hours post-stimulation (**Fig. 1c**). Two-way ANOVA revealed a significant main effect of KCl treatment, time point and interaction between the two for *Bdnf I, II, IV, VI* and *IX* transcripts. Sidak’s multiple comparisons post-hoc test showed that *Bdnf I* and *IV* are significantly increased 3 hours post-stimulation (*p* < 0.0001), which was also reflected by significant upregulation of total *Bdnf IX* at this time point (*p* < 0.0001) (**Fig. 1c**). Interestingly, *Bdnf V* did not respond to KCl stimulation at any time point as indicated by the lack of main effect of KCl treatment, time point, or interaction (*p* = 0.7595). Additionally, *Bdnf III* expression was significantly decreased at all time points, as indicated by the significant main effect of KCl treatment (p < 0.0001), but not time point (p = 0.6056). We did not detect transcript variants *VII* and *VIII* [10, 26]at baseline or after KCl stimulation (data not shown), and therefore these transcripts were not measured in subsequent experiments.

**Figure 1.**
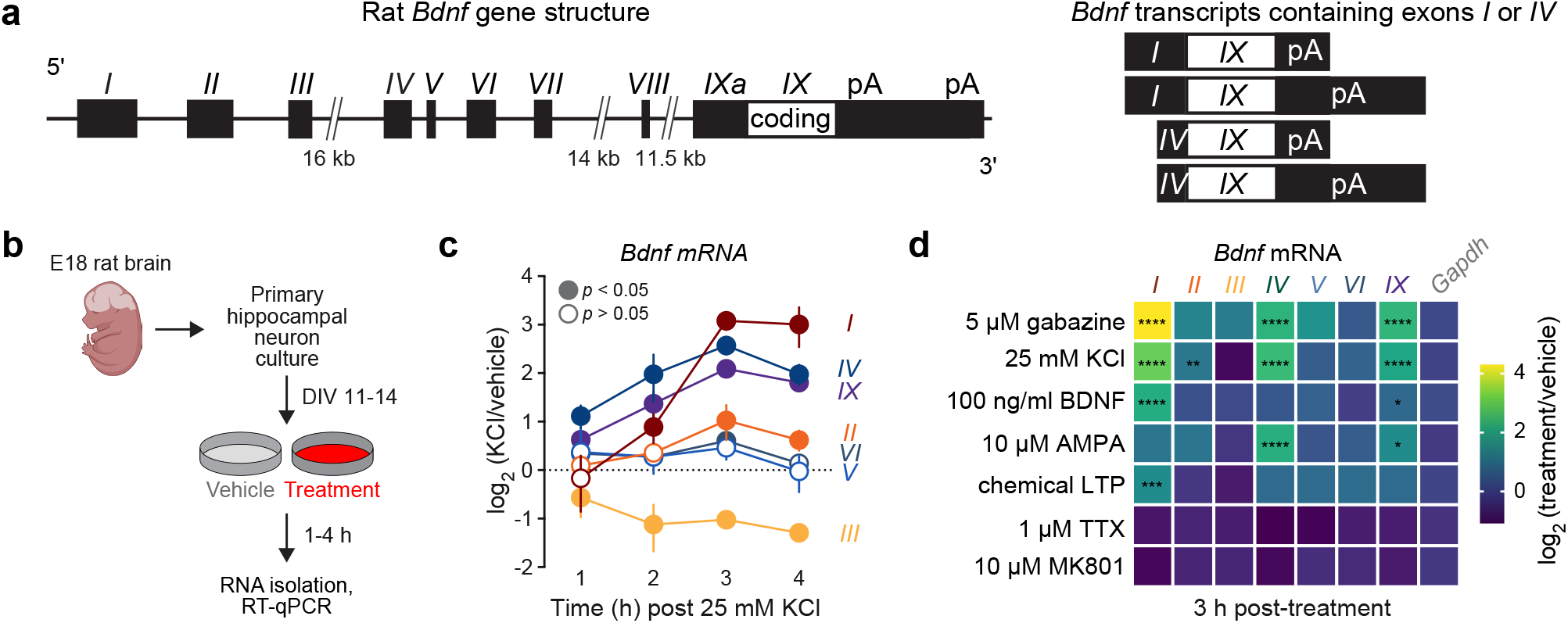
*Bdnf* transcripts *I* and *IV* are most responsive to activity. **a**, *Bdnf* gene structure illustrating non-coding exons (*I-IXa*) and common-coding exon (*IX*) (left). Schematic of the *Bdnf* transcript variants containing non-coding exons *I* or *IV* spliced to the common-coding exon *IX* with short or long 3’ untranslated regions (UTRs) (right). **b**, Rat hippocampal neuron culture preparation. **c**, Expression of *Bdnf I* and *IV* transcript variants, as well as total *Bdnf IX*, peaks 3 h after depolarizing rat hippocampal cultures with 25 mM KCl (n = 6, two-way ANOVA; 3 h *Bdnf I* p < 0.0001; 3 h *Bdnf IV* p < 0.0001; 3. *Bdnf IX* p < 0.0001; Sidak’s multiple comparisons test). Expression of other *Bdnf* transcripts is up-or downregulated at different time points after KCl (n = 6, two-way ANOVA; 3 h. *Bdnf II* p = 0.0141; 3 h *Bdnf III* p < 0.0001; 3 h *Bdnf VI* p = 0.0185; Sidak’s multiple comparisons test). **d**, Stimulation of rat hippocampal cultures with 5 μM Gabazine, 25 mM KCl, 100 ng/ml recombinant BDNF, chemical LTP (200 nM NMDA; 50 μM forskolin; 0.1 μM rolipram), and 10 μM AMPA for 3 h upregulated the expression of select *Bdnf* transcript variants, with *Bdnf I* and *Bdnf IV* transcripts being most responsive (n = 6-12, one-way ANOVA; 5 μM gabazine F(7, 48) = 118.8, p < 0.0001; 100 ng/ml BDNF F(7, 88) = 99.66, p < 0.0001; 10 μM AMPA F(7, 48) = 9.947, p < 0.0001; chemical LTP F(7, 48) = 6.594, p < 0.0001). Blockade of neuronal activity with 1 μM tetrodotoxin (TTX) or 10 μM MK801 for 3 h. inhibited *Bdnf* expression, affecting mostly *Bdnf I* and *Bdnf IV* transcripts (n = 8, one-way ANOVA; 1 μM TTX F(7, 56) = 1.142, p = 0.3505; 10 μM MK801 F(7, 56) = 1.461, p = 0.2002). Data in panel c are expressed as mean ± SD. Individual comparisons; *p < 0.05, **p < 0.01, ***p < 0.001, ****p < 0.0001.

Because activity-dependent changes in *Bdnf* transcription vary in response to neuronal stimulation and inhibition protocols [10, 11, 13], we quantified the expression of *Bdnf* variants in response to multiple treatment protocols (**Fig. 1d**). DIV 11 rat hippocampal neuron cultures were treated with the GABA_A_ receptor antagonist gabazine (5 μM), KCl (25 mM), recombinant BDNF (100 ng/ml), AMPA (10 μM), the sodium channel blocker TTX (1 μM), and the NMDA receptor antagonist MK801 (10 μM), as well as a chemical LTP protocol (200 nM NMDA, 50 μM forskolin and 0.1 μM rolipram). RNA was isolated 3 hours after stimulation, and RT-qPCR revealed that patterns of *Bdnf* transcript expression were unique for different treatment protocols. For example, gabazine stimulation produced the strongest upregulation of total *Bdnf IX* levels (5.5-fold), with a 20-fold upregulation of *Bdnf I* and 5-fold upregulation of *Bdnf IV* (**Fig. 1d**). While recombinant BDNF and chemical LTP treatments only upregulated *Bdnf I* 4.6-fold and 2.8-fold, respectively, AMPA stimulation upregulated *Bdnf IV* by 5-fold. These results demonstrate that the pattern of stimulus-dependent *Bdnf* transcript variant expression is unique to different environmental stimuli and that *Bdnf* variants *I* and *IV* are most highly regulated by activity.

### CRISPRa targeting of *Bdnf I* or *Bdnf IV* drives expression of the respective transcript and corresponding BDNF protein production

We recently developed a CRISPR activation (CRISPRa) system that successfully targets *Bdnf I* and *Bdnf IV* transcripts [20]. We deployed this system to better understand whether *Bdnf* transcripts possess unique biological functions (**Fig. 2a**). Single guide RNAs (sgRNAs) were designed complementary to 19-20 nucleotide sequences within *Bdnf* promoters *I* and *IV* less than 50 base pairs (bp) upstream from the transcription start sites (TSS) (**Fig. 2b**; **Table S1**). dCas9-VPR and sgRNAs were packaged into high-titer (10^11^ GC/ml) lentiviral vectors and co-transduced into primary rat hippocampal cultures at DIV 4-5 (**Fig. 2c**). Robust transgene expression was verified by immunocytochemistry (ICC) with antibodies specific for mCherry (to visualize sgRNA) and FLAG (to visualize dCas9-VPR) at DIV 11 (**Fig. S1**).

**Figure 2.**
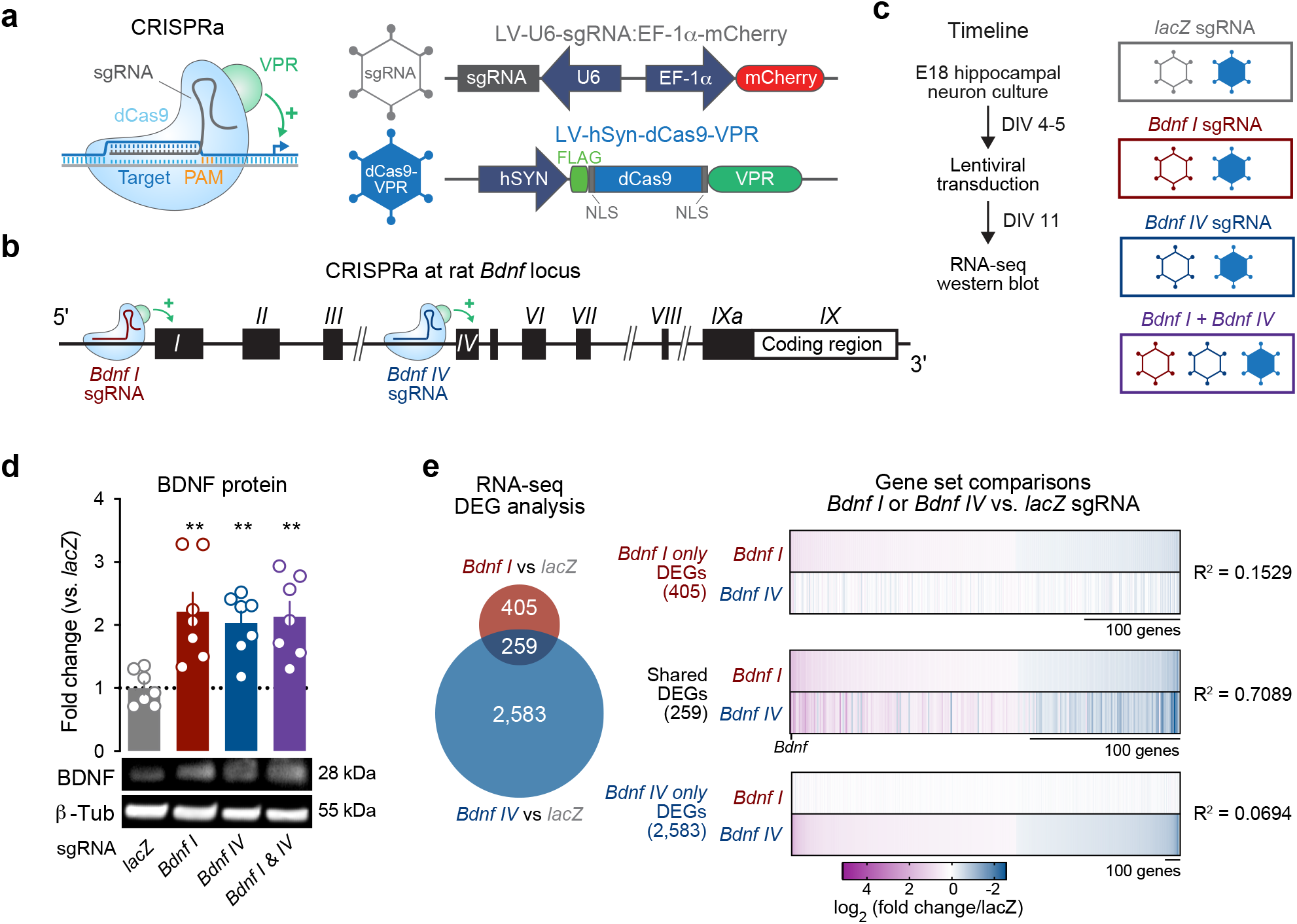
CRISPRa induction of *Bdnf I* or *IV* transcript variants upregulates BDNF protein levels and causes distinct changes in downstream gene expression. **a**, Schematic of the CRISPR/dCas9-VPR system and associated plasmids. **b**, Schematic of CRISPRa targeting of *Bdnf* promoters *I* and *IV*. **c**, Experimental timeline and viral transduction groups. **d**, Western blot for BDNF protein after CRISPRa targeting of *Bdnf I, Bdnf IV*, and *Bdnf I & IV* transcripts revealed a significant increase in BDNF protein levels in all conditions as compared to the non-targeting *lacZ* control (n = 7, one-way ANOVA, F(3, 24) = 6.696, p = 0.0019; Dunnett’s multiple comparisons test). **e**, Venn diagram illustrating 664 differentially expressed genes (DEGs) from RNA-seq after *Bdnf I* sgRNA targeting (red) and 2,842 DEGs after *Bdnf IV* sgRNA targeting (blue), as compared to *lacZ* sgRNA control. Right, correspondence heatmaps indicating the overlap between DEGs after *Bdnf I* upregulation and *Bdnf IV* upregulation. Simple linear regression analysis indicated that expression changes between manipulations were at least partially overlapping (significant derivation from zero, p < 0.0001 for all comparisons), with varying effect sizes (indicated by R^2^ values). All data are expressed as mean ± SEM. Individual comparisons; **p < 0.01.

We previously confirmed selective upregulation of *Bdnf* transcripts *I* or *IV* with virtually no off-target CRISPRa effects [20]. CRISPRa produced a twofold increase in mature BDNF protein levels after either *Bdnf I, Bdnf IV* or *Bdnf I & IV* targeting as compared to a non-targeting negative control, *lacZ* gRNA (**Fig. 2d**). These data suggest that upregulation of either *Bdnf* transcript is equally capable of producing mature BDNF protein.

### *Bdnf I* or *Bdnf IV* upregulation with CRISPRa causes unique gene expression changes

BDNF plays a key role in regulating synaptic plasticity by modulating downstream signaling cascades [11, 27, 28]. However, whether manipulation of distinct *Bdnf* transcripts differentially impacts downstream gene expression profiles has not been explored. We previously reported that CRISPRa of *Bdnf I* or *Bdnf IV* in hippocampal neurons resulted in 259 common differentially expressed genes (DEGs) when compared to a *lacZ* control (**Fig. 2e**) [20]. In agreement with *Bdnf*’s role in activity-dependent signaling cascades, we found both that *Bdnf I* and *Bdnf IV* upregulation resulted in regulation of immediate early genes (*Fos, Arc, Egr1* and *Egr3*). However, we identified many DEGs that were unique to CRISPRa conditions - 405 DEGs specific to *Bdnf I* upregulation and 2,583 DEGs specific to *Bdnf IV* upregulation, compared to *lacZ* control (**Fig. 2e**) [20]. To determine whether these DEGs are unique to the upregulation of each *Bdnf* transcript or whether there is a correspondence between DEGs in one condition and the same genes in the other condition, we directly compared log2 fold change values of *Bdnf I* only DEGs to the same genes in the *Bdnf IV* condition (**Fig. 2e**, top heatmap) and, vice versa, *Bdnf IV* only DEGs to the same transcripts in the *Bdnf I* sgRNA condition (**Fig. 2e**, bottom heatmap). Linear regression of log2 fold change values between conditions revealed that there was a low (yet significant, *p* < 0.0001) correlation between *Bdnf I* and *Bdnf IV* manipulations. These data indicate that independent upregulation of either *Bdnf I* or *Bdnf IV* produces both unique as well as overlapping sets of gene expression changes.

### DEGs in *Bdnf I* vs. *Bdnf IV* CRISPRa conditions are associated with unique gene ontology terms and neuropsychiatric disorders

Correspondence analysis (**Fig. 2e**) indicated at least partial overlap between *Bdnf I* or *Bdnf IV* CRISPRa targeting. However, closer inspection revealed that many genes were also oppositely regulated by these manipulations, suggestive of distinct transcriptional consequences of isoform-specific upregulation. To more definitively identify transcripts that were uniquely regulated following either *Bdnf I* or *Bdnf IV* CRISPRa, we next performed a new DESeq2 analysis to directly compare these two CRISPRa-targeting conditions (*Bdnf I* vs. *Bdnf IV* sgRNAs). This analysis revealed 1,483 genes selectively enriched after *Bdnf I* upregulation and 1,943 genes selectively enriched after *Bdnf IV* upregulation (**Fig. 3a**; **Tables S2, S3**). Gene ontology (GO) analysis identified some functional overlap between *Bdnf I* vs. *IV* - regulated genes, such as neuron differentiation, axon development, neuron projection development, and mRNA processing (**Fig. 3b**), suggesting that these cellular processes depend on expression of both *Bdnf I* and *IV* transcripts. However, each set of *Bdnf I* vs. *IV* - regulated genes was also associated with unique GO terms. *Bdnf I*-regulated genes belonged to GO categories associated with transcription, dendrite development, neurogenesis and differentiation of specific brain areas and cell types (**Fig. 3b**, top). *Bdnf I*-regulated genes were responsible for dendrite development, such as *Shank1, Shank2, Ntrk2*, and *Nrep* [29], and activity-dependent gene expression, such as *Mecp2* [30] (**Table S4**).

**Figure 3.**
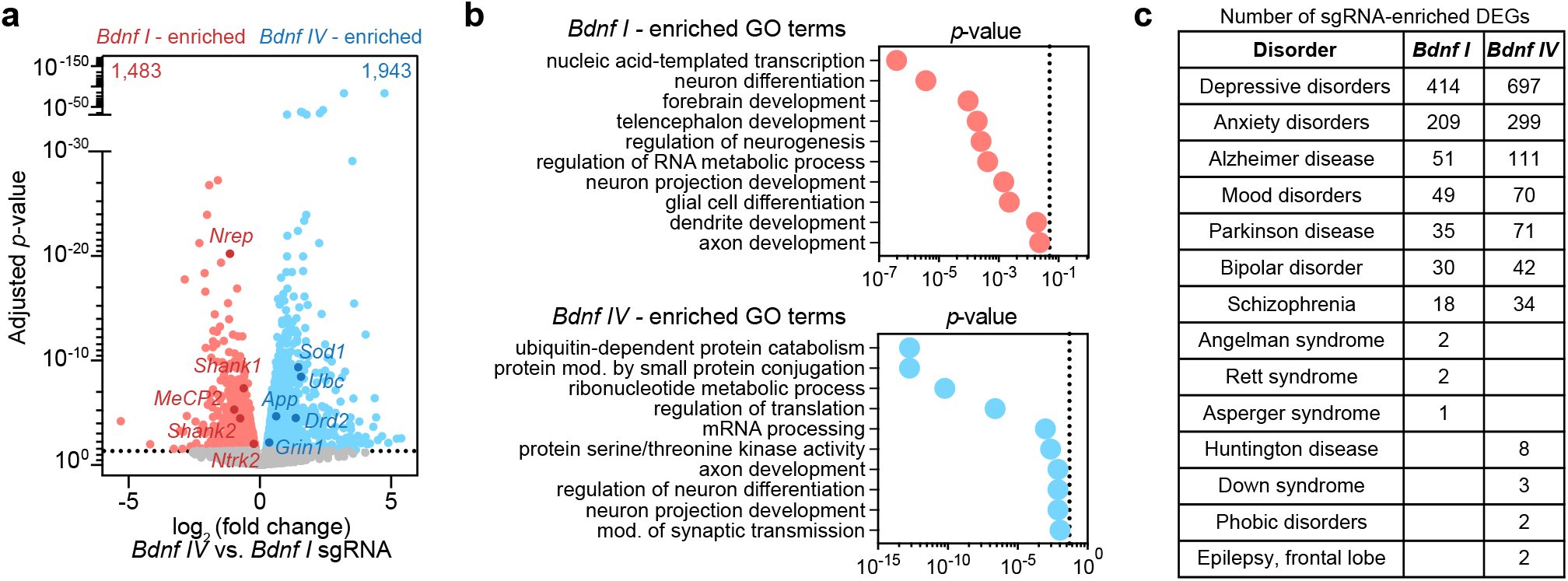
RNA-seq directly comparing *Bdnf I* vs. *Bdnf IV* sgRNA-targeting with CRISPRa revealed unique DEGs associated with different gene ontology terms and neuropsychiatric disorders. **a**, Volcano plot showing DEGs detected by DESeq2 in dCas9-VPR *Bdnf IV* vs. *Bdnf I* - targeted conditions. Standard cutoff point is represented by the horizontal dotted line (adjusted p < 0.05). *Bdnf I* - enriched (1,483 genes, red) and *Bdnf IV* - enriched (1,943 genes, blue) genes are indicated. **b**, Top significant gene ontology (GO) terms for *Bdnf I* - enriched and *Bdnf IV*- enriched genes, illustrating some overlapping (axon development, neuron projection development, neuron differentiation) but mostly unique GO terms for each gene set. **c**, DEGs after *Bdnf I* or *Bdnf IV* upregulation with CRISPRa overlapping clinical gene sets in the Harmonizome database.

*Bdnf IV*-regulated genes were associated with key cellular processes such as protein catabolism and metabolism, and regulation of translation (**Fig. 3b**, bottom). The top GO terms associated with *Bdnf IV* upregulation were ubiquitin-dependent protein catabolism and modification of proteins by small protein conjugation, such as ubiquitination, with specific enriched genes like *Ubc* and other ubiquitin-proteasome pathway-related genes (**Table S5**). *Bdnf IV*-regulated genes were known synaptic plasticity and neurodegeneration-associated marker genes, such as *App, Sod1, Grin1*, and *Drd2* (**Fig. 3a,b**). These data imply that upregulation of distinct *Bdnf* transcripts may be associated with specific intracellular pathways that have physiological significance for cell-type and brain-area specific processes.

*Bdnf* expression is frequently altered in neurocognitive and neuropsychiatric disorders [31, 32]. Therefore, we investigated whether identified DEGs after *Bdnf I* vs. *Bdnf IV* upregulation overlapped with MESH term-defined clinical gene sets within the Harmonizome database (Harmonizome_CTD Gene-Disease Associated Dataset) [33]. We identified that genes differentially expressed after *Bdnf I* and *Bdnf IV* upregulation were associated with psychiatric disorders, including depression, anxiety, bipolar disorder, schizophrenia and mood disorders, as well as neurocognitive diseases, such as Alzheimer’s and Parkinson’s (**Fig. 3c**). In addition, we found that some neurological disorders were uniquely associated with DEGs in either *Bdnf I* or *Bdnf IV*-manipulated conditions. For example, Angelman, Rett and Asperger syndromes were uniquely associated with genes differentially expressed after *Bdnf I* upregulation (**Table S6**), while Huntington’s disease, Down syndrome, phobic disorders and frontal lobe epilepsy were associated with DEGs after *Bdnf IV* upregulation (**Table S7**) (**Fig. 3c**). While these neurocognitive disorders may have shared gene dysregulation profiles, unique *Bdnf* transcripts may have differential roles in disease etiology.

### *Bdnf I* upregulation with CRISPRa increases mushroom dendritic spine density, length, and head diameter, and increases the complexity of dendritic arbors

*Bdnf* enhances dendritic branch length and number [3, 34, 35] and promotes synaptic spine formation and maturation [2, 36], while disrupting BDNF expression decreases spine density [37]. Previous studies reported that selective disruption of BDNF production from distinct promoters has opposing effects on CA1 and CA3 dendritic arbors and dendritic spine morphology [38]. Therefore, we investigated whether CRISPRa upregulation of *Bdnf I* and *Bdnf IV* contributes to dendritic spine density and arborization.

Rat hippocampal primary cultures were transduced with CRISPRa constructs on DIV 4-5 and transfected with a construct encoding Lifeact-GFP on DIV 12, a fluorescently-tagged small peptide which binds to intracellular f-actin [39] (**Fig. 4a**). Cells were fixed at DIV 14 and 63X high-resolution confocal laser scanning microscopy z-stack images were acquired for dendritic spine analysis. After maximum-intensity image deconvolution, three-dimensional digital reconstruction models of dendrites were quantified based on length and shape (**Fig. 4b**). Mushroom spine density was significantly increased after *Bdnf I* and *Bdnf I & IV* upregulation, as compared to a *lacZ* control (**Fig. 4c**, mushroom). Furthermore, mushroom spine volume, length and spine head diameter were increased specifically after *Bdnf I*, but not *Bdnf IV*, upregulation (**Fig. 4d**).

**Figure 4.**
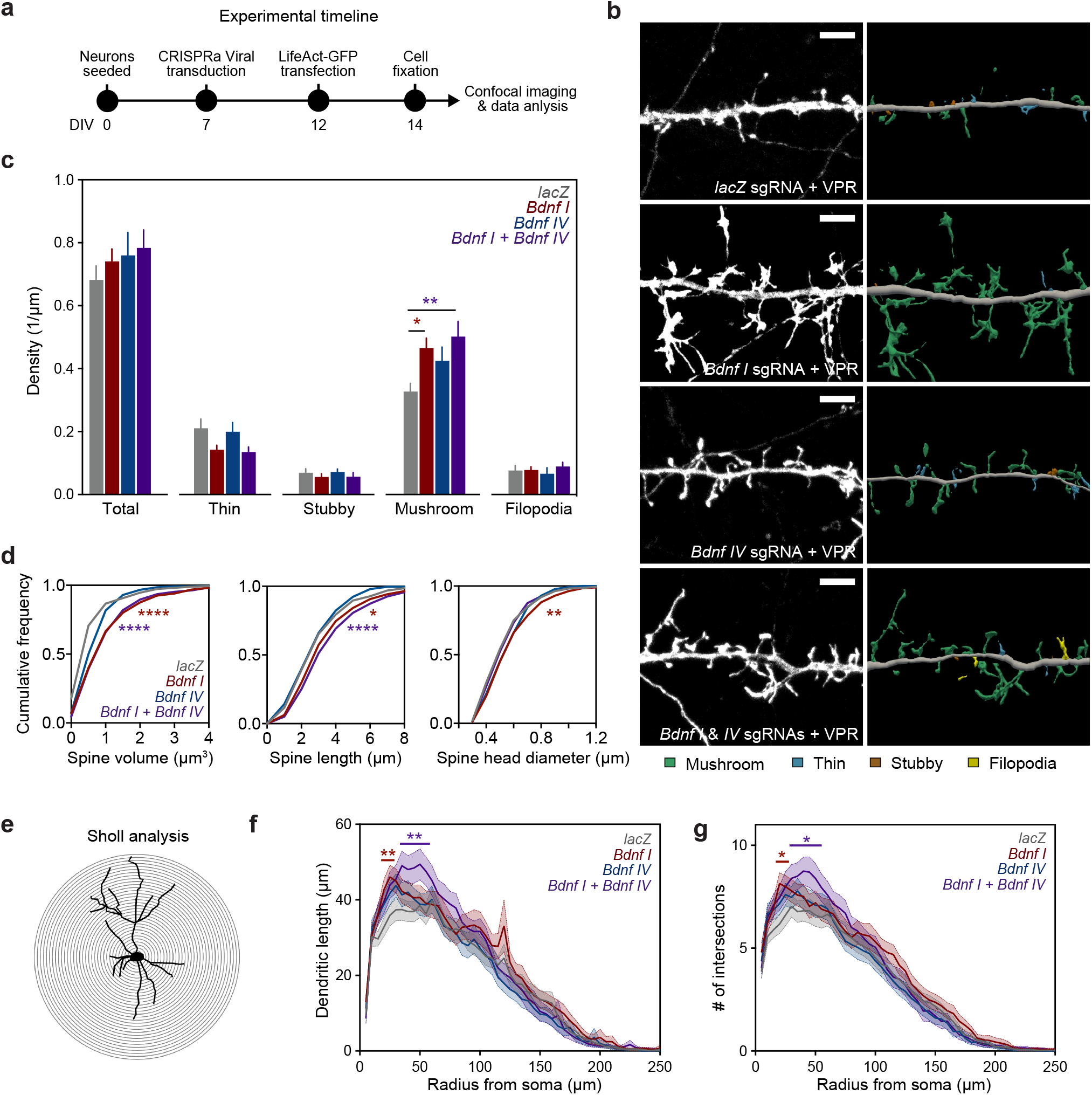
Mushroom spine density, volume, length and head diameter as well as dendritic complexity are significantly increased after *Bdnf I*, but not *Bdnf IV*, upregulation. **a**, Experimental design timeline. **b**, Representative maximum-intensity high-resolution confocal microscope images after deconvolution (left) and corresponding three-dimensional digital reconstruction models of dendrites (right) after *Bdnf I, Bdnf IV* or *Bdnf I & IV* targeting with dCas9-VPR. Scale bar, 5 μm. Colors in digital reconstructions correspond to dendritic classes: green represents mushroom spines; blue, thin spines; brown, stubby spines; and yellow, filopodia. **c**, Dendritic spine density of thin, stubby, mushroom and filopodia spines, as well as all combined spines (total), illustrating a significant increase in mushroom spine density after CRISPRa targeting of *Bdnf I* and *Bdnf I & IV*, compared to a *lacZ* control (n = 15-24 dendrites from three independent cell cultures, one-way ANOVA, F(3, 71)= 3.878, p = 0.0126). **d**, The cumulative frequency distributions for mushroom spine volume (left), length (center), and head diameter (right) plotted for each *Bdnf* or *lacZ* dCas9-VPR-targeting condition. *Bdnf I* and *Bdnf I & IV* upregulation with CRISPRa significantly increases mushroom spine volume and length (n = 225-388 spines, one-way ANOVA, F(3, 1223) = 16.42, p < 0.0001) while Bdnf I upregulation significantly increases mushroom head diameter (n = 225-388 spines, one-way ANOVA, F(3, 1223) = 5.127, p = 0.0016). **e**, Schematic of the Sholl analysis using concentric circles starting at the soma in 5 μm increments (outlined by Lifeact-GFP fluorescence). **f**, Dendritic length in each concentric segment, as a function of the radius from the soma, illustrating that CRISPRa upregulation of *Bdnf I* and *Bdnf I & IV* significantly increases proximal dendrite length (n = 20-29 dendritic branches from three individual cultures, two-way ANOVA, F(50,4800) = 220.7, p < 0.0001). **g**, The number of dendritic intersections as a function of the radius from the soma, illustrating that CRISPRa upregulation of *Bdnf I* and *Bdnf I & IV* significantly increases the number of proximal dendritic intersections (n = 20-29 dendritic branches from three individual cultures, two-way ANOVA, F(49,4704) = 271.2, p < 0.0001). All data are expressed as mean ± s.e.m. Individual comparisons, *p < 0.05, **p< 0.01, ****p < 0.0001.

To explore whether upregulation of *Bdnf I* or *IV* influences dendrite complexity, rat hippocampal primary neurons were transduced with CRISPRa constructs and transfected with Lifeact-GFP (**Fig. 4b**). Cells were fixed at DIV 14 and 20X high-resolution confocal laser scanning microscopy z-stack images were acquired. Neuronal 3D reconstructions for Sholl morphometry analyses with concentric circles were performed (**Fig. 4e**). CRISPRa upregulation of *Bdnf I*, but not *Bdnf IV*, significantly increased dendritic length (**Fig. 4f**) and the number of dendritic intersections at the 25-75 μm radius from the cell body, as compared to a *lacZ* control (**Fig. 4g**). These data illustrate that *Bdnf I* upregulation increases the length and complexity of dendritic arbors.

### Upregulation of *Bdnf IV*, but not *I*, decreases freezing during contextual fear learning

BDNF plays a critical role in many forms of learning and memory, including the acquisition and extinction of contextual fear memories [40–43]. Therefore, we next investigated whether *Bdnf I* and *IV* differentially impact contextual fear learning. We bilaterally infused high-titer (10^12^ GC/ml) lentiviruses expressing dCas9-VPR and sgRNAs for either *Bdnf I, IV, I & IV* or *lacZ* into the CA1 subregion of the dorsal hippocampus of adult rats (**Fig. 5a**). After a 12-day recovery period to allow for the stable integration and expression of the virally-delivered transgenes (**Fig. 5b**), rats underwent contextual fear conditioning (CFC) (**Fig. 5c**) as previously described [25]. Rats were handled for two days immediately prior to training and then were exposed to the training chamber for 2 min, followed by three 1 sec, 0.5 mA shocks delivered every 2 min, with a final exploration period of 1 min after the final shock. Rats were re-exposed to the same fear conditioning chamber 1 hour, 24 hours, and 7 days after the training for 7 min and time spent freezing was recorded.

**Figure 5.**
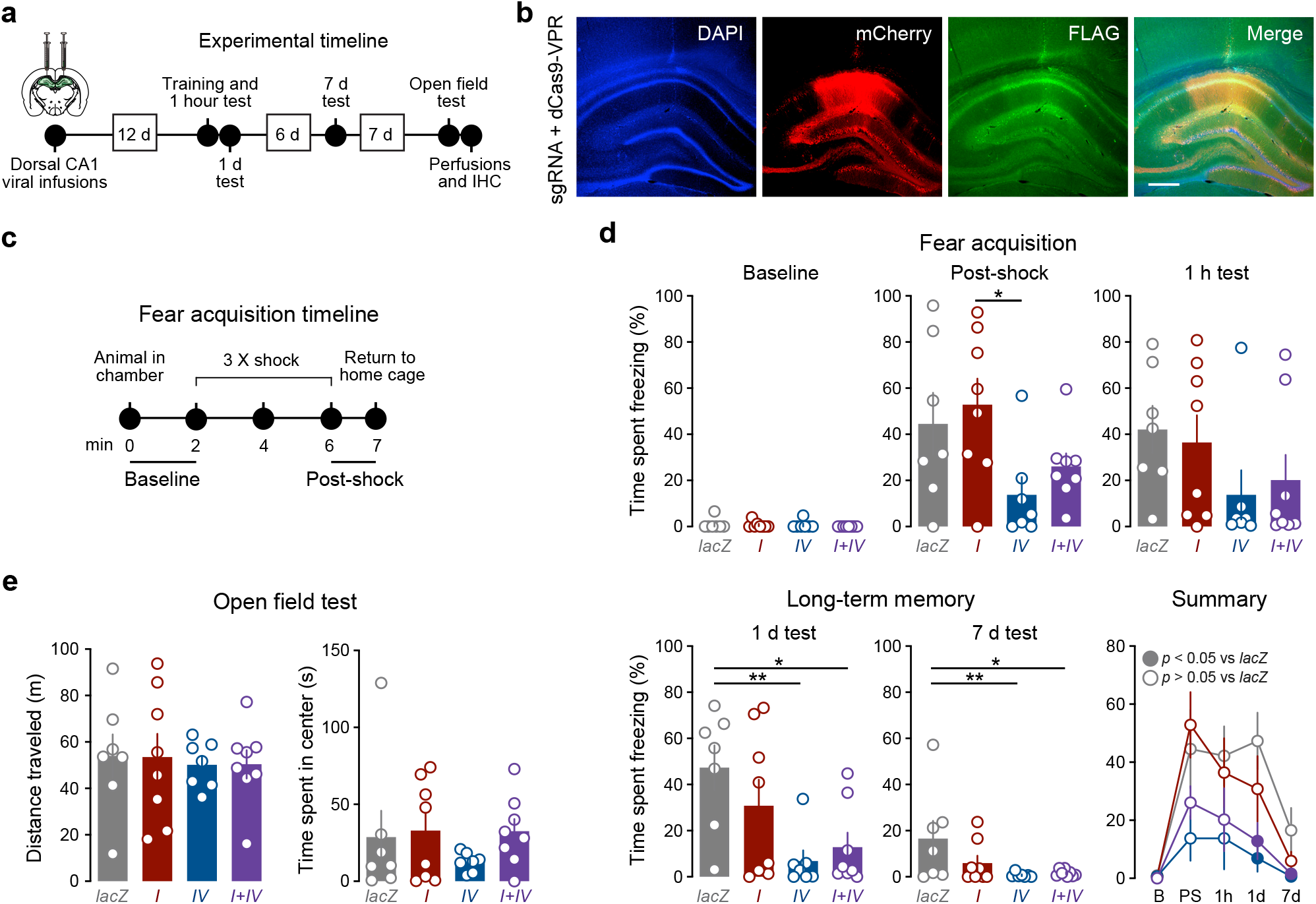
CRISPRa-mediated induction of *Bdnf IV*, but not *Bdnf I*, decreases time spent freezing following contextual fear conditioning (CFC). **a-b**, Bilateral lentiviral infusions targeting dorsal CA1 of the hippocampus were performed in adult male rats (n = 7-8 rats/condition). Two weeks after stereotaxic viral infusions, CFC and open field tests were carried out, followed by transcardial perfusions and immunohistochemistry (IHC) to verify sgRNA (mCherry, red) and dCas-VPR (FLAG, green) expression. Cell nuclei were stained with DAPI. Scale bar, 500 μm. Schematic of the target region was adapted from Paxinos and Watson. **c**, Schematic of fear acquisition (training) timeline. **d**, Comparison of freezing behavior at different timepoints before and after CFC. There were no significant differences between groups at baseline (prior to CFC). Animals in the *Bdnf IV* CRISRPa condition exhibited a significant decrease in time spent freezing as compared to *Bdnf I* CRISPRa condition when tested immediately after CFC (n = 7-8, one-way ANOVA; F(3,26) = 3.948). While no significant differences were observed one hour following CFC, *Bdnf IV* and *Bdnf I & IV* CRISPRa rats exhibited significantly lower time spent freezing when tested 1 d or 7 d after CFC compared to the *lacZ* controls (n = 7-8, one-way ANOVA; 1 d test: F(3,26) = 4.421; 7 d test: F(3,26) = 3.169). All data are expressed as mean ± s.e.m. Individual comparisons, *p < 0.05 and **p < 0.01. **e**, There were no significant differences in the total distance traveled or time spent in the center of the arena in the open field test.

Rats in all conditions had low baseline levels of freezing prior to shock presentation (**Fig. 5d**, left), and time spent freezing increased following training. Rats in the *Bdnf IV*-upregulated condition spent significantly less time freezing compared to rats in the *Bdnf I*-upregulated condition during the last minute of training after the final shock (**Fig. 5d**). While not statistically significant, *Bdnf IV*-upregulated rats spent less time freezing compared to *lacZ* and *Bdnf I* rats, when tested 1 hour after conditioning (**Fig. 5d**). When tested at 24 hours and 7 days, there was a significant reduction in time spent freezing in the *Bdnf IV*, as well as *Bdnf I & IV*-upregulated conditions (**Fig. 5d**). To rule out that changes in time spent freezing were due to changes in activity or anxiety, we quantified total locomotion and center time in an open field test 7 days following the last fear conditioning test. There were no significant differences in total distance traveled or time spent in the center of the arena between any of the *Bdnf*-targeting conditions or the *lacZ* control (**Fig. 5e**).

Taken together, these data suggest that upregulation of *Bdnf IV*, but not *I*, decreases expression of contextual fear.

## DISCUSSION

Prior efforts to study the role of BDNF in the brain includes infusion of recombinant BDNF [41, 44–46], region- and cell type-specific conditional knockdowns [16, 47], and mouse models lacking *Bdnf* production from distinct promoters [48]. To expand our understanding of *Bdnf* gene regulation in neuronal plasticity and behavior, we developed and validated tools for precise manipulation of transcript-specific *Bdnf* expression using CRISPRa/i from the endogenous loci [20, 49]. Using CRISPRa, we investigated distinct roles of activity-dependent *Bdnf* transcripts *I* and *IV* in cultured hippocampal neurons and in the rat dorsal hippocampus. Our findings *in vitro* suggest that overexpressing *Bdnf I* vs *IV* has differential effects on downstream gene expression as well as on dendrite length and dendritic spine morphology. *In vivo*, we find that overexpressing *Bdnf I* vs *IV* differentially impacts fear expression in response to CFC.

While upregulation of *Bdnf I* and *IV* similarly increased BDNF protein levels (**Fig. 2d**), there were differential effects on gene expression (**Fig. 2e**). These results support the possibility that each splice variant is associated with unique downstream molecular signaling pathways. *Bdnf I*-regulated genes were associated with dendrite development, neurogenesis and differentiation of specific brain areas and cell types (**Fig. 3b**, top), results that support the finding that mushroom spine phenotypes (increased densities, spine volume, length and head diameter) were increased and that more complex dendritic arbors developed in response to *Bdnf I*, but not *Bdnf IV*, upregulation (**Fig. 4c-g**). Moreover, genes enriched after *Bdnf I* upregulation in the “dendrite development” GO category (including *Cdkl5, Mecp2, Disc1, Shank1, Shank2*, and *Ntrk2*) (**Table S4**) were also identified within the Harmonizome clinical gene set database to be associated with Angelman, Rett and Asperger syndromes (**Fig. 3c**), which are associated with dendrite and dendritic spine dysgenesis [50, 51]. These findings provide novel insights on the role of *Bdnf I* dendritic spine development in neurodevelopmental disease regulation.

In contrast, *Bdnf IV*-regulated genes belonged to GO categories that control basic cellular functions, such as protein catabolism and metabolism (**Fig. 3b**, bottom). Interestingly, *Bdnf IV* upregulation was strongly associated with enrichment of genes encoding proteins in the ubiquitin-proteasome pathway (UPP), which mediates regulated degradation of intracellular proteins [52, 53]. Genes encoding ubiquitin, *Ubc*, proteasomal alpha subunits, *Psma1-6*, proteasomal beta subunits, *Psmb3, 4, 7, 9*, and proteasomal ATPases, *Psmc2-5*, were enriched after *Bdnf IV* upregulation (**Table S5**). The UPP is necessary for induction of the transcription and translation-dependent late phase of LTP (L-LTP), as well as learning and memory consolidation [53, 54], and chemically blocking the UPP prevents induction of L-LTP and *Bdnf* transcription [55, 56]. While further investigation is needed, it is plausible that upregulating *Bdnf IV* triggers an autoregulatory loop engaging further transcription of UPP components and enhanced UPP activity.

Both *Bdnf I* and *Bdnf IV*-regulated genes were associated with similar neuropsychiatric and neurodegenerative disorders, including depression, anxiety, Alzheimer’s and Parkinson’s diseases, disorders where a relationship with *Bdnf* dysregulation is well-documented [57, 58]. However, only *Bdnf IV*-regulated genes were associated with Huntington’s disease (*App, Drd2*), Down syndrome (*Sod1*), phobic disorders (*Casp3*), and frontal lobe epilepsy (*Chrna4, Chrnb2*), also previously tied to *Bdnf* dysregulation [31, 59]. Together with the RNA-seq data, these findings suggest that although the downstream transcriptional signatures caused by *Bdnf I* and *Bdnf IV* upregulation are similar, there are many unique downstream transcriptional readouts associated with each splice variant that may be important for neuronal function and disease progression.

While the importance of *Bdnf* expression in CFC has been well established, contributions of individual *Bdnf* transcripts is not clear. Mutant mice in which production of BDNF from promoter *IV* is disrupted show enhanced fear and resistance to fear extinction [42], which can be rescued by selective over-expression of BDNF in hippocampal neurons that project to the prefrontal cortex [43]. Supporting these previous studies, our data show that upregulating *Bdnf IV*, but not *I*, in dorsal CA1 of the rat decreases fear expression 24 hours and 7 days after CFC. In our study, *Bdnf IV*-targeted animals trended towards lower freezing levels even 1 hour after fear conditioning, and hence further studies are necessary to disentangle whether the decreases in fear expression at later time points are due to a general decrease in expression of fear, deficits in fear memory acquisition, or enhanced fear extinction. Further studies deciphering the role of the *Bdnf IV* transcript in the regulation of learning and memory and CFC are necessary and, are now possible, due to the novel CRISPR tools described here and in previous manuscripts [49, 60–62].

In summary, our study is the first to report the functional impacts of manipulating different transcript variants of *Bdnf* both in rat hippocampal neurons and in rat hippocampus. CRISPRa upregulation of *Bdnf I* promotes formation of mature dendritic spines and extended dendritic arbors, while upregulation of *Bdnf IV* mediates expression of contextual fear. Future studies delineating distinct roles for *Bdnf* transcript variants in the nervous system will increase our understanding of *Bdnf’*s involvement in synaptic plasticity, neuropsychiatric and neurodegenerative diseases, as well as our ability to modulate *Bdnf* gene expression for therapeutic applications.

## ACKNOWLEDGEMENTS

We thank all current and former Day Lab members for assistance and support. L.I. is supported by the Civitan International Research Center at UAB. We acknowledge support from the University of Alabama at Birmingham Biological Data Science Core (RRID:SCR_021766), the UAB Heflin Center for Genomic Sciences.

## AUTHOR CONTRIBUTIONS

SVB:Conceptualization,Methodology,Formal Analysis, Investigation, Writing - Original Draft, Writing - Review & Editing, Project Administration, Funding Acquisition

AJB: Investigation

DH: Investigation

JJT: Investigation

LI: Formal analysis, Data Curation

KMG: Investigation

BWH: Investigation

JHH: Resources

KM: Writing - Review & Editing, Supervision

JJD: Conceptualization, Writing - Review & Editing, Supervision, Project Administration, Funding Acquisition

## SUPPLEMENTAL METHODS

### Animals

All experiments were approved by the University of Alabama at Birmingham Institutional Animal Care and Use Committee (IACUC). Male Sprague Dawley rats (90- to 120-day old and weighing 250-350 g) and timed pregnant dams were purchased from Charles River Laboratories. Dams were individually housed until embryonic day (E)18 for hippocampal cell culture harvest. Male rats were co-housed in pairs in plastic cages in an IACUC-approved animal care facility on a 12/12 h light/dark cycle with *ad libitum* food and water. Animals were randomly assigned to experimental conditions.

### Neuronal cell cultures

E18 rat hippocampal tissue was used to generate primary rat hippocampal cultures, as described previously [20, 21]. Briefly, cell culture plates (Denville Scientific Inc.) were pre-coated with poly-L-lysine (Sigma-Aldrich; 50 μg/ml) supplemented with 7.5 μg/ ml laminin (Sigma-Aldrich) overnight and rinsed twice with diH2O the following day. Hippocampal tissue from E18 rat pups was digested with papain (Worthington LK003178) for 25 min at 37°C and rinsed in complete Neurobasal media (supplemented with B27 and L-glutamine, Invitrogen). Sequential trituration through three fire-polished Pasteur pipettes (large to small pipette mouth openings) was used to generate a single cell suspension. Cells were filtered through a 100-μm cell strainer (Fisher Scientific), pelleted, resuspended in fresh media, counted, and seeded to a density of 125,000 cells per well on 24-well culture plates (65,000 cells/cm^2^). For immunocytochemistry and dendritic spine experiments, cells were grown on glass coverslips (12 mm; Carolina Biological Supply). Cells were maintained in complete Neurobasal media for 11 - 14 d *in vitro* (DIV) in a humidified CO2 (5%) incubator at 37°C with half media changes at DIV1, DIV4–DIV5, DIV8–DIV9, and DIV12. Upon completion of each experiment, the media was aspirated from cell culture plates and the plates were either frozen on dry-ice for gene expression studies or fixed with 4 % paraformaldehyde (Sigma) for staining.

### Lifeact-GFP transfection

Plasmid encoding Lifeact-GFP was a generous gift from Dr. Gary Bassell, Emory University. Neurons were transfected at DIV12 with Lifeact-GFP using Lipofectamine 2000 (Invitrogen), according to manufacturer instructions. At DIV14, cells were fixed with 4% PFA, washed with 1x PBS, and coverslips were mounted onto glass sliced (Fisher, Catalog #12-550-15) using Vectashield mounting media (Vector La Catalog #H1000). Z-series images were acquired on a Zeiss LSM-800 confocal microscope with a 60X oil immersion objective at 0.15μm increments through the entire visible dendrite. Dendrites selected for imaging were secondary branches, a minimum of 25 μm away from the soma, and non-overlapping with other dendrites. Prior to analysis, captured images were deconvolved using Huygens Deconvolution System (16.05, Scientific Volume Imaging, the Netherlands) with the following settings: CMLE; maximum iterations: 50; signal to noise ratio: 40; quality: 0.01.

### Cell culture chemical stimulation

DIV11 hippocampal cell cultures were subjected to a variety of chemical treatment protocols. 25 mM KCl reconstituted in complete cell culture media was applied for 1 - 4 hours. 5 μM Gabazine (Sigma-Aldrich), 100 ng/ml recombinant BDNF (PeproTech), 10 μM AMPA (Sigma-Aldrich), 1 μM tetrodotoxin (Sigma-Aldrich), and 10 μM MK801 (Sigma-Aldrich) were reconstituted in complete cell culture media and applied for 3 hours. Chemical LTP treatment consisted of sequential application of 200 nM NMDA (Millipore-Sigma) for 10 min, followed by 50 μM forskolin (Millipore-Sigma) and 0.1 μM rolipram (Millipore-Sigma) for 15 min, as previously described [63]. NMDA, forskolin and rolipram were reconstituted in complete cell culture media. Vehicle-treated cell cultures received fresh media at corresponding time points. After chemical stimulation, media was aspirated and cells were frozen on dry ice and stored at -80°C for later RNA extraction.

### RNA extraction and RT-qPCR

Total RNA was extracted using an RNeasy kit (QIAGEN) and reverse-transcribed using iScript cDNA Synthesis kit (Bio-Rad) per manufacturer’s guidelines. cDNA was subject to RT-qPCR for genes of interest, as described previously [20]. A list of qPCR primer sequences is provided in **Table S1**. Forward *Bdnf* primers were designed complementary to each non-coding exon and reverse primer complementary to the common-coding *Bdnf* exon *IX*. To determine the combined expression level of all *Bdnf* transcripts, both forward and reverse primers were designed within the common-coding *Bdnf IX*.

### CRISPR/dCas9 and sgRNA construct design

A lentivirus compatible backbone (a gift from Feng Zhang, Addgene #52961; [64]) was used to insert a dCas9-VPR (VP64-p65-Rta) cassette driven by the human synapsin 1 promoter (SYN1) [20]. SP-dCas9-VPR was a gift from George Church (Addgene #63798; [19]). A guide RNA scaffold (a gift from Charles Gersbach, Addgene #47108; [65]) was inserted into a lentivirus compatible backbone containing EF1α-mCherry. A BbsI cut site within the mCherry construct was mutated with a site-directed mutagenesis kit (NEB). *Bdnf*-specific sgRNA targets were reused from our previous study [20] and were designed using CHOPCHOP (http://chopchop.cbu.uib.no/; [66]). All CRISPR RNA (crRNA) sequences were analyzed with National Center for Biotechnology Information’s (NCBI) Basic Local Alignment Search Tool (BLAST) to ensure specificity. An extensive validation of minimal sgRNA off-target effects was conducted in our previous study [20]. The bacterial *lacZ* gene was used as a non-targeting sgRNA control.

### Lentivirus production

Lentiviruses were produced in a sterile environment subject to BSL-2 safety. HEK-293T cells (ATCC catalog # CRL-3216) were grown in T225 culture flasks in standard HEK media (DMEM + heat-deactivated 10% FBS) and passaged at 70% confluence. For viral production, 100% confluent HEK-293T cells were co-transfected with the dCas9-VPR or the sgRNA plasmid, along with the psPAX2 packaging plasmid, and the pCMV-VSV-G envelope plasmid (Addgene #12260; Addgene #8454, respectively) using FuGene HD transfection reagent (Promega) for 40–48 h in HEK cell medium. Supernatant medium was passed through a 0.45 μm filter and centrifuged at 25,000 rpm for 1 h 45 min at 4°C in a refrigerated floor ultracentrifuge (Bechman Coulter). The viral pellet was resuspended in 100 μl of sterile 1X PBS, and stored at –80°C in single use aliquots. Physical viral titer was determined using Lenti-X RT-qPCR Titration kit (Takara), and only viruses > 1 × 10^10^ GC/ml or > 1 × 10^12^ GC/ml were used for cell culture or *in vivo* experiments, respectively.

### Western blot

Total protein was precipitated from the first flow-through an RNeasy mini column (QIAGEN). Each protein sample was resuspended in RIPA lysis buffer (50 mM Tris-HCl, 150 mM NaCl, 1% NP-40, 0.5% sodium deoxycholate, 0.1% SDS, and 1X Halt protease and phosphatase inhibitor; Pierce), boiled at 95°C for 5 min with 4X Laemmli buffer (Bio-Rad), separated on a 4–15% polyacrylamide gel, and transferred to a polyvinylidene difluoride membrane. BDNF was detected with a rabbit monoclonal anti-BDNF primary antibody (1:1000; Abcam; ab108319) and a goat anti-rabbit secondary antibody (1:10,000; IR dye 800, LI-COR Biosciences; 827-08365). Blots were visualized on an Azure c600 imaging system (Azure Biosystems). β-Tubulin was detected using a mouse anti-β-Tubulin antibody (1:2000; Millipore; 05-661) with a goat anti-mouse secondary antibody (1:10,000; IR dye 680, LI-COR Biosciences; 926-68170), as a loading control. Protein levels were quantified in ImageJ, and BDNF intensity values were normalized to β-Tubulin. For rat neuronal BDNF quantification, proBDNF (∼28 kDa) appeared as the dominant BDNF signal over mature BDNF (∼13 kDa), and was used for quantification. Recombinant BDNF (Peprotech; 450-02-10UG) was used as a positive control.

### Immunocytochemistry and immunohistochemistry

Immunocytochemistry (ICC) was performed as described previously [20]. To validate expression of the dCas9-VPR or the sgRNA plasmids, cells cultured on glass coverslips (12 mm; Carolina Biological Supply) were fixed with 4% paraformaldehyde (Sigma), blocked with 10% Thermo Blocker BSA and 1% goat serum, and incubated in an anti-FLAG primary antibody (1:5000 in 1X PBS; Thermo Fisher Scientific; MA1-91878) or an anti-mCherry primary antibody (1:200 in 1X PBS; Thermo Fisher Scientific; PA5-34974) overnight at 4°C. Fluorescent secondary antibodies, Alexa Fluor 488 goat anti-mouse and Alexa Fluor 555 goat anti-rabbit (1:500; Thermo Fisher Scientific; A-10667 and A-78954, respectively), were applied the following day for one hour at room temperature in the dark. Cells on coverslips were mounted onto microscope slides (Fisher Scientific) with Prolong Gold antifade medium (Invitrogen) containing 4,6-diamidino-2-phenylindole (DAPI) as a marker for cell nuclei. For immunohistochemistry (IHC), adult male rats were transcardially perfused with formalin (1:10 dilution in 1X PBS, Fisher). Brains were removed and postfixed for 24 h in formalin, then sliced at 50 μm using a vibratome. Brain slices were permeabilized with 0.25% Triton X-100 in 1X PBS and the IHC steps performed as for ICC described above. 63X images were taken on a Zeiss LSM-800 confocal microscope.

### Contextual fear conditioning

Two weeks following stereotaxic surgery, animals were habituated to handling for two consecutive days by the experimenters and contextual fear conditioning (CFC) was carried out the next day. CFC was conducted as previously described [25]. Animals were transported and kept outside of the behavioral suite for at least 2 h prior to experimentation. Rats were placed into the training chamber with a metal floor grid (Med Associates) and allowed to explore for 2 min (baseline freezing), after which they received three electric shocks (1 s, 0.5 mA each) every 2 min. After final shock, animals were allowed to explore the context for an additional 1 min (freezing after final shock) before being returned to their homecage. Memory performance was tested at 1 h, 24 h, and 7 days after training by returning the animals to the conditioned chamber and observing freezing behavior using high-speed video recording (Med Associates). CFC data from animals with no viral expression or incorrect hippocampal dorsal CA1 targeting (as assessed by IHC) were excluded from data analysis.

### RNA-sequencing (RNA-Seq) and data analysis

RNA-Seq was conducted at the University of Alabama at Birmingham Heflin Center for Genomic Core Laboratories. RNA was extracted, purified (RNeasy, QIAGEN), and DNase-treated for three biological replicates per experimental condition. RNA was prepared for directional sequencing using SureSelect Strand Specific RNA Library Prep kit (Agilent Technologies) per manufacturer’s recommendations. Poly A RNA libraries underwent sequencing (75-bp paired-end directional reads; 22–38 M reads/sample) on an Illumina sequencing platform (NextSeq2000). Paired-end FASTQ files were uploaded to the University of Alabama at Birmingham’s High Performance Computer cluster for custom bioinformatics analysis using a pipeline built with snakemake, as previously described [20]. Differentially expressed genes (DEGs) were designated if they passed a *p* < 0.05 adjusted *p* value cutoff and contained basemeans > 50. Venn diagrams and Volcano plot were constructed in Prism (GraphPad). DEGs were compared against MESH term-defined clinical gene sets in the Harmonizome database [33]. Gene ontology (GO) analysis was conducted with co-regulated genes (genes either up- or down-regulated by *Bdnf I* or *Bdnf IV* sgRNA treatments, as compared to *LacZ* sgRNA control) using the Cytoscape software with a ClueGO plug in [67].

### Stereotaxic surgery

Naïve adult Sprague Dawley rats were anesthetized with 4% isoflurane and secured in a stereotaxic apparatus (Kopf Instruments) under aseptic conditions. An anesthetic plane was maintained with 1–2.5% isoflurane throughout the surgical procedure. Guide holes were drilled using stereotaxic coordinates for the dorsal CA1 region of the hippocampus (AP: –3.3 mm, ML: ±2.0 mm from bregma). All infusions were made using a gastight 30-gauge stainless steel injection needle (Hamilton Syringes) that extended –3.1 mm (DV from bregma) into the infusion site. Bilateral lentivirus micro infusions of 1.5 μl (0.5 μl sgRNA and 1 μl dCas9-VPR viruses in sterile 1X PBS) per hemisphere were made using a syringe pump (Harvard Apparatus) at a rate of 0.25 μl/min. Injection needles remained in place for 10 min following the infusion to allow for viral diffusion. Guide holes were covered with sterile bone wax and surgical incision sites were closed with nylon sutures. Animals received buprenorphine and carprofen for pain management and topical bacitracin to prevent infection at the incision site. Animals were allowed to recover from surgery for two weeks prior to behavioral training.

### Statistical analysis

Statistical and graphical analyses were conducted with Prism (GraphPad Software, La Jolla, CA). Data are presented as mean ± SEM. All graph error bars represent SEM. Statistical significance was designated at 0.05 for all analyses. One-way ANOVA with Dunnett’s multiple comparisons test or two-way ANOVA with Sidak’s multiple comparisons test were used where appropriate. To compare spine densities between experimental conditions, the mean spine density was calculated per experimental replicate. These experiment means were then averaged per experimental condition.

### Data availability

Sequencing data that support the findings of this study have been deposited in Gene Expression Omnibus (GEO) with the accession number GSE117961. CRISPRa constructs have been deposited, along with plasmid maps and sequences, in the Addgene plasmid repository (RRID:Addgene_114196; RRID:Addgene_114199).

---

**Figure S1.**
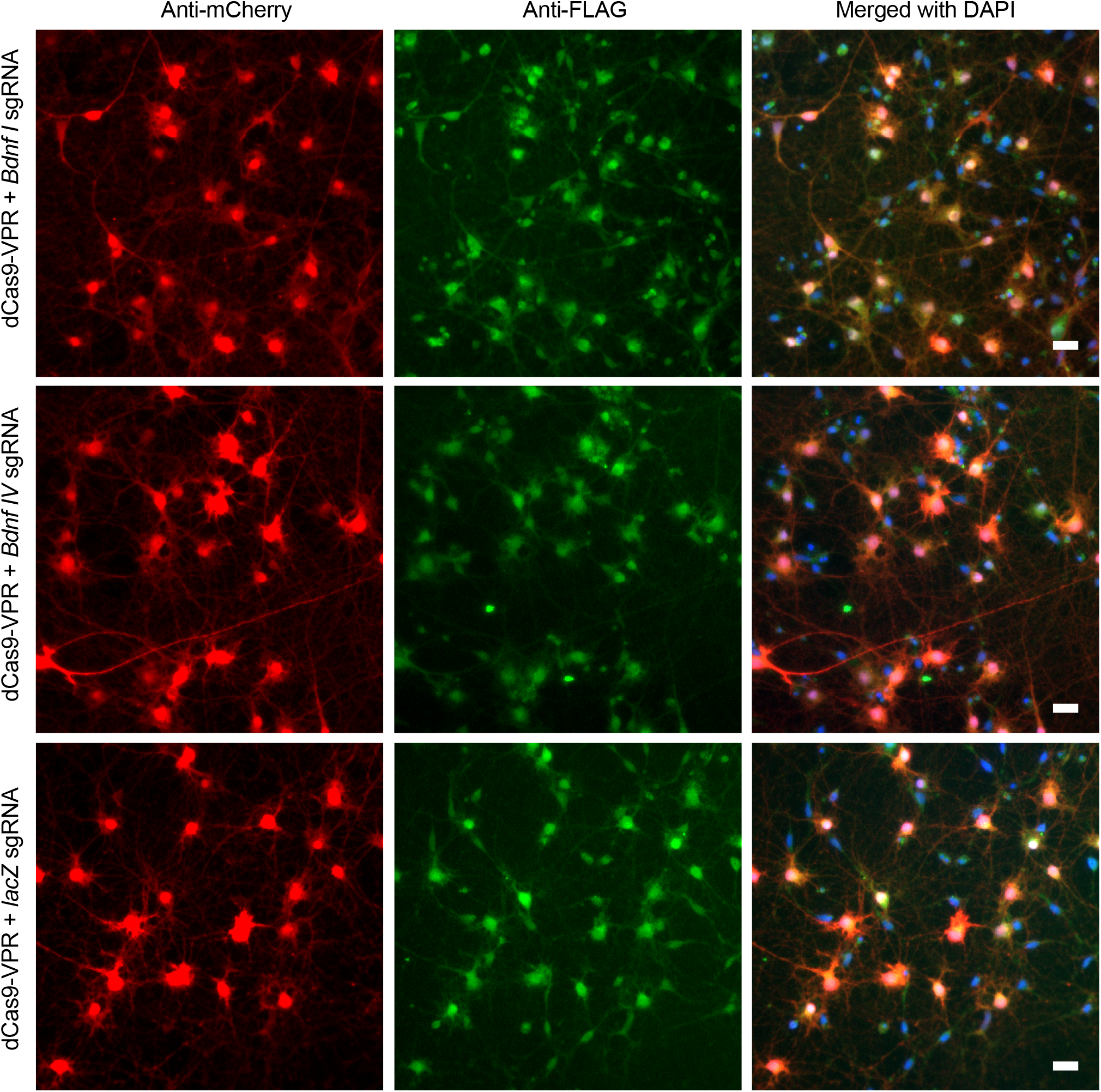
Verification of successful co-transductiondCas9-VPR and sgRNAs in rat primary hippocampal neurons. Antibodies against mCherry (to visualize sgRNAs, red) and FLAG (to visualize dCas9-VPR, green) were used for immunocytochemistry. High level of overlap between mCherry and FLAG signals in the merged image (white) confirm successful co-transduction of both constructs in cell cultures. Scale bar, 20 μm.

